# Effects of repetitive Iodine Thyroid Blocking on the Development of the Foetal Brain and Thyroid in rats: a Systems Biology approach

**DOI:** 10.1101/710764

**Authors:** David P.A. Cohen, Mohamed Amine Benadjaoud, Phillipe Lestaevel, Dalila Lebsir, Marc Benderitter, Maâmar Souidi

## Abstract

A single dose of potassium iodide (KI) against repeated exposure to radioactive iodine, such as the one of the Fukushima accident, might not be effective enough to protect the thyroid. Our group showed that repetitive dose of KI for eight days offers efficient protection without toxic effects in adult rats. However, the effect of repetitive KI on the developing foetus still unknown especially on brain development, but a correlation between the impaired maternal thyroid status and a decrease in intelligence quotient has been observed. In this study, gene expression analysis of the progeny of repetitive KI-administered pregnant rats performed by our group showed distinct gene expression profile from two different organs: thyroid and cortex. To understand how these differentially expressed genes are implicated in the observed behaviour change, a systems biology approach was used to construct networks using three different techniques; Bayesian statistics using ShrinkNet, sPLS-DA on the DIABLO platform using mixOmics and manual construction of a Process Descriptive network. For each organ, we were able to construct gene expression network, to select genes that are most contributing to either control or KI-treated groups, respectively, and to construct the PD network from differentially expressed (DE) gene enriched with data from publications. Furthermore, we were able to connect DE genes from both organs into one network with genes from both organ participating in the same cellular processes that affect mitophagy and neuronal outgrowth.

This work may help to evaluate the doctrine for using KI in case of repetitive or prolonged exposure to radioactive particles upon nuclear accidents.

## 1 INTRODUCTION

During a nuclear emergency, large quantities of radioactive particles may be released in a plume contaminating the environment and populations likely to be exposed. Isotopes present in the discharges include radioactive iodine-131 (^131^I). Exposure of populations to radioactive iodine may be responsible in the absence of appropriate protective measures for the increase in risk for late-onset thyroid cancers, particularly in children, such as been observed in the territories most affected by the fallout from the Chernobyl accident^1^ or the Hiroshima and Nagasaki bombing^2^.

To prevent health consequences, protective actions during the early phase of a nuclear emergency are recommended, including population sheltering, Iodine Thyroid Blocking (ITB) provision, evacuation and restrictions on the consumption of water or agricultural products^3^. The objective of this ITB is to saturate the thyroid gland with non-radioactive iodine and thus avoid the fixation of radioactive iodine (Wolff-Chaikoff effect)^4^.

However, the recent 2011 Fukushima Daiichi disaster showed that World Health Organization’s “ITB guidelines”^3^ that so far supports a single intake of potassium iodide (KI) tablets cannot adequately protect populations in case of prolonged (beyond 24 hours) or repeated exposure. Indeed, the “ITB guidelines” has been based on the assumption that populations would be exposed relatively shortly to radioactive discharges from which they would be rapidly subtracted. In a situation of prolonged and/or repeated discharges, a second intake of stable iodine is possible, although the “ITB guidelines” so far give no indication about repeated administration for the implementation KI^3,5^. However, the repetitive ITB recommendation by the WHO is not necessarily the optimal implementation for sensitive sub-populations. Therefore more knowledge about new stable iodine administrations is essential for effective prevention of thyroid pathologies.

Recently, preclinical data, obtained by our group, on adult rats, have showed that a daily administration during several days of stable iodine does not induce adverse outcome (AO) ^6,7^. However, after ITB for several consecutive days in gestational rats, certain adverse effects on the central nervous system have been observed on the offspring^8,9^. We hypothesized in this article that transient hypothyroidism caused by the dose regimen for repeated ITB in pregnant rats, leads to congenital hypothyroidism in the foetus and consequently to neurotoxicity.

In this study, an *in utero* rat model has been subjected to repetitive KI administration over eight days during the brain and thyroid development of the foetus. A system biology approach was proposed to identify a putative molecular mechanism that explains the effect of KI on the thyroid and cortex. To associate genes with the AO impacting the Central Nervous system (CNS) observed by Lebsir and colleagues^8^, transcriptomics was applied to identify genes that are significantly differentially expressed (DE) in the thyroid and the cortex, respectively. From these genes, correlation between them was inferred using two different methods. To determine a putative mode of action, a Process Descriptive (PD) diagram has been constructed using the analysed DE genes and their correlation data enriched with data obtained from the PubMed database. The PD diagram represents molecular mechanisms^10^ and identifies genes belonging to key events that may be associated with the AO. In addition, identification of causal relationships between genes from the thyroid and the cortex was explored.

## 2 MATERIALS AND METHODS

### Protocol experiments

Animals were housed in the IRSN animal facilities accredited by the French Ministry of Agriculture for performing experiments on rodents. Animal experiments were performed in compliance with the French and European regulations on protection of animals used for scientific purposes (EC Directive 2010/63/EU and French Decree 2013–118). All experiments were approved by the Ethics Committee #81 and authorized by the French Ministry of Research under the reference APAFIS#10827-201707312156679 v2 (internal project number P14-06).

The study included sixteen pregnant Wistar rats (Charles River laboratories, L’arbresle, France), divided into control group receiving NaCl solution, and treated group receiving KI 1 mg/kg over eight days since the 9th gestational day (GD9 – GD16). The NaCl (pH 7.4) and KI (1 mg/kg) solutions were prepared and kindly provided by the Central Pharmacy of Armed forces (Orleans, France).

They were housed individually upon arrival and allowed to recover from transportation for one week. Rats were kept in regular light/dark schedule (12h/12h), at 21 ± 2°C and 50 ± 10% humidity. Food 0.3 mg I / kg of pellet (A04-10 SAFE, Augy, France) and water were freely accessible. After birth and weaning, male pups were randomly separated from their mother almost one pup from each mother to avoid inbreeding bias. After that, they were divided into control progeny not exposed *in utero* to KI, and treated progeny exposed *in utero* to KI, each group include fourteen animals. Thirty days after the weaning, thyroid and cerebral cortex progeny were collected and instantly deep-frozen in liquid nitrogen and then stored at −80°C.

### Transcriptomics

Transcriptomics on thyroid and cortex was routinely performed by CRIBIOM (Marseille, France) using the One-colour Microarrays-Based gene expression analysis, low input Quick Amp labelling version 6.9.1 protocol (Agilent). An amount of 100 ng of total RNA was hybridised with Cy-3 and subsequently hybridised on microarrays (SurePrint G3 8×60K, Agilent). The raw fluorescence signals were scanned with DNA Microarray Scanner SureSelect (Agilent) and analysed with the R-package Limma^11^.

### Network inference

The Gene network inference was performed using the R-package Shrinknet v1.0 with parameters suggested by the authors^12^ using expression data obtained from transcriptomics. ShrinkNet calculates the interaction (edges) between genes/nodes and a set of edges defines the topology of a network which may generate useful hypothesis about the adverse effect of repetitive ITB. Only edges with a FDR lower than 0.05 were kept to construct the network.

### sPLS-DA and DIABLO framework

Besides the previous gene network inference approach, a supervised multivariate analysis was conducted by means of the R package mixOmics (version 6.6.1) in R studio^13^ to specifically target inter-group differences (KI and Controls). sPLS-DA enables the selection of the most predictive or discriminative features in the data to classify the samples^14^. sPLS-DA helps to prioritise genes of interest for the construction of the PD network.

Firstly, two sparse partial least square discriminant analysis (sPLS-DA) models were conducted separately on cortex and thyroid transcriptomics’ result. After this intra organ analysis, an inter organ transcriptomics integration was conducted via the DIABLO framework^15^. DIABLO constructs components across the cortex and thyroid transcriptomics matrices, maximizing their covariance given a control or KI treatment group as a response variable. These components are defined as linear combinations of transcriptomics variables and a L1 penalization ensures their sparsity. In all the analyses (inter and intra organ), Leave-One-Out Cross validation was used to determine the model parameters including number of components and number of features per component.

The results were carried out in order to obtain a relevance network (Correlation threshold = 0.82) using the “network” command and permits the selected features representation and highlight positive or negative association in the within/between organ connections.

The cytoscape software^16^ v.3.420 was used to visualize the networks obtained by all the network inference approaches (Shrinknet, sPLS-DA and DIABLO).

### PD Network construction

The manual construction of the biochemical network depicting thyroid hormone biosynthesis and iodide metabolism started from the collection of common genes highlighted simultaneously by all the methods presented in the network inference section above (Shrinknet, sPLS-DA and DIABLO), assuming that it constitutes the most robust significant gene candidates. Relevant information from scientific articles was retrieved from the PubMed database using the molecule(s) of interest as key word(s), e.g. “gene A” was used as a query or the combination either “thyroid” or “cortex”, respectively, and “gene A” was used as query. “Gene B” was used as separate query either in combination with key words or not. No data-mining tools were used. The CellDesigner software (version 4.4)^17^ was used to represent molecular biological mechanisms, derived from the literature, resulting in a structured network representation (PD diagram) compliant with Systems Biology Markup Language (SBML) level 2 that is suitable for further computational analysis^18^. Each reaction has been annotated in the “reaction note” at least once with a corresponding scientific article. The CellDesigner’s graphical notation^19^ can be converted into the Systems Biology Graphical Notation (SBGN) standard^20^ by using the CellDesigner’s internal convertor. For each PD diagram, the CellDesigner software creates an xml-file that can be downloaded from the supplementary material section. In addition the PD diagrams are exported to a svg-file format that can be opened in any internet browser and the diagram can viewed with different zoom levels. These files can be downloaded from the supplementary material section as well.

### Colouring molecules in the PD network according to gene expression

The genes that are differentially regulated (transcriptomic analysis) and are present in the PD network can be coloured according to their log2 FC values by using the Cytoscape plugin BiNoM^21^ with Cytoscape v2.8. With the BiNoM plugin the CellDesigner file can be loaded into Cytoscape environment and colours can be assigned to genes and proteins according to their fold change.

## 3 RESULTS

### Gene expression profile for the thyroid and cortex

Transcriptomic analysis revealed 60 and 94 annotated transcripts for the thyroid and cortex, respectively (t-test, Q-value < 0.05) (Figure 1). Certain functions could be associated to these genes including site-specific DNA binding, ATP binding, transcription from the thyroid and transcription regulation, DNA, RNA and ATP binding, nucleotide binding and neurogenesis for the cortex (Table 1 in the supplementary materials). These lists of transcripts or genes were used to construct a network to attempt finding a molecular mechanism. The heat map showed a distinct difference in gene expression between the control and the KI-exposed group.

**Figure 1.**
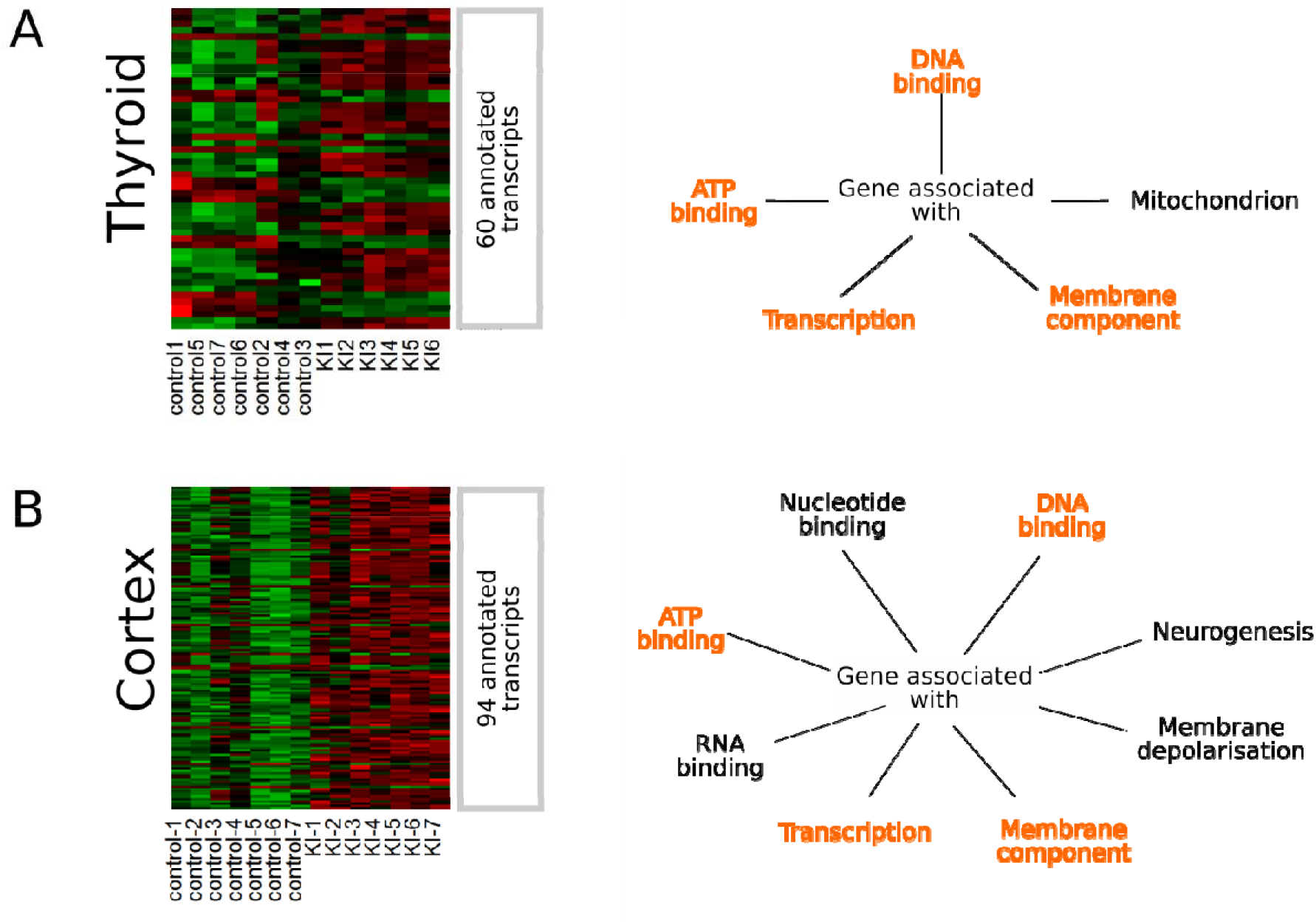
Heat map of significant differentially expressed (DE) genes and their associated function or cellular localisation isolated from progeny rats after 30 days of weaning in the thyroid (1A) and the cortex (1B). The progeny have been divided into two groups; one group exposed *in* utero to KI treatment while the control group was not exposed to KI in *utero*. Figure 1A shows a heat map of DE genes and some of their associated function and cellular localisation of genes obtained from the thyroid. Figure 1B shows a heat map of DE genes and some of their associated function and cellular localisation of genes obtained from the cortex.

### Network inference for the thyroid upon repetitive KI treatment

Bayesian statistics showed that the 60 transcripts/genes from the thyroid can be connected into four major sub-networks and few genes (nodes) that connect to many other nodes, many nodes that connect to a few nodes (Figure 2A).

**Figure 2.**
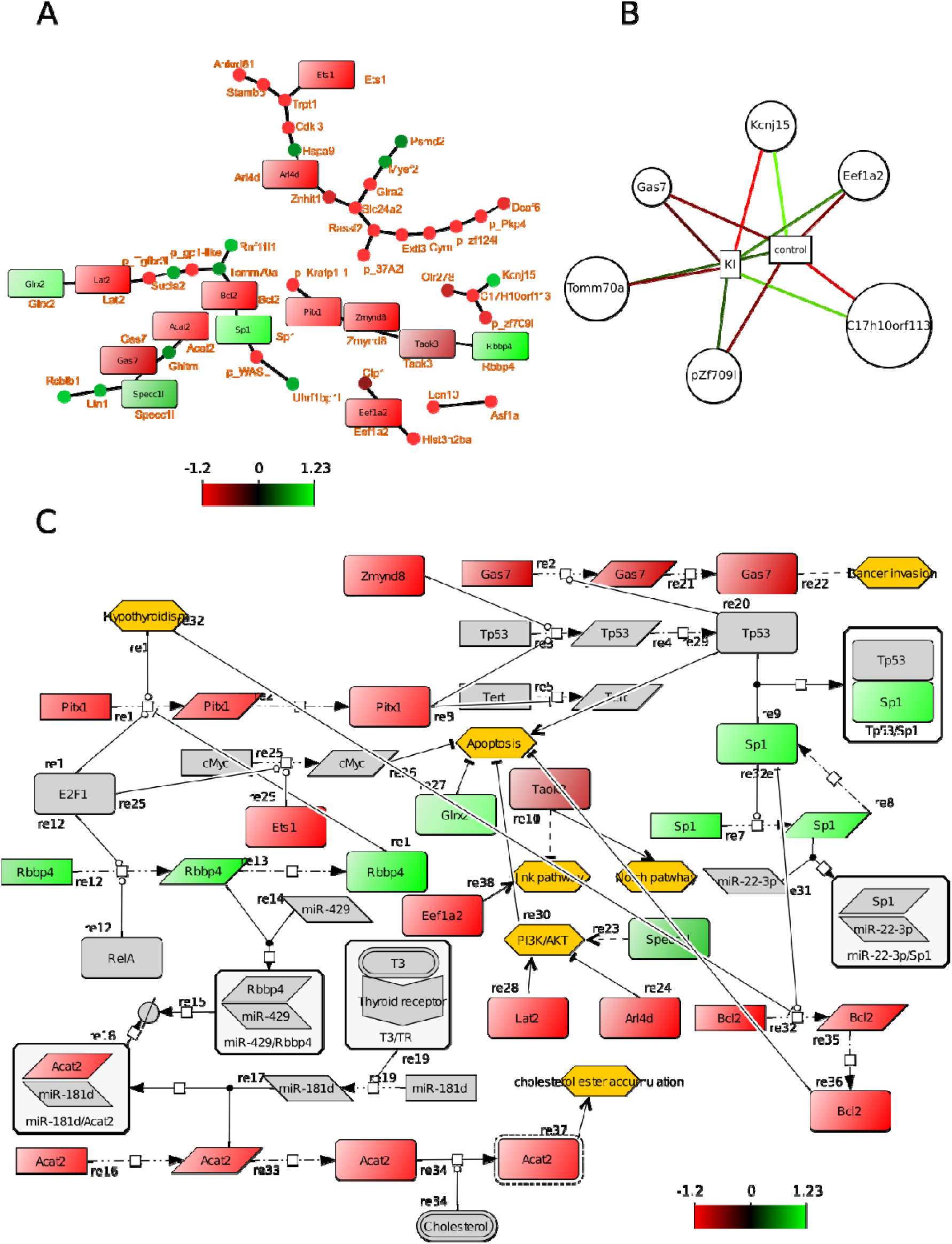
Selected diagrams show correlations between genes from the thyroid. The colour of the nodes indicates up or down regulation. Ochre coloured hexagons show phenotypes/biological processes that are modulated by the network. Figure 2A shows a undirected network of DE genes using Bayesian simultaneous-equation model with global-local shrinkage^12^. The nodes (closed circles and squares) represent genes and edges represent correlation between genes. The square nodes are genes that can be found in the process descriptive network as well (figure 2C). Figure 2B shows the results of sPLS-DA: genes that contribute most to either the control group or the IK exposed group, respectively. These genes will be used as a starting point for constructing the PD network (figure 2C). Figure 2C shows a process descriptive network that represents molecular mechanism^10^. Squares that are colour coded shows the fold changes of DE genes. The grey shapes are molecules obtained from published articles. All edges have been described before in the scientific articles.

This inferred network gave indications for constructing a PD network e.g. which proteins might interact with other molecules. How many different un-related processes might be present? The colour code indicated a log2 fold change of the genes in the KI-treated group compared to the control. Genes for the constructing the PD network were prioritised: sPLS-DA method was applied to obtain genes that contribute to the different phenotypes (control and KI exposed groups). All genes selected by sPLS-DA except for *tomm70a* and *kcnj15* were under expressed in the KI-exposed group (Figure 2B). The combination of the results of Bayesian statistics and sPLS-DA was used as input for constructing the PD network. This network contained 45 chemical species including genes, RNA protein, anti-sense RNA molecules and 37 edges that have been annotated (Figure 2C). The annotations can be found back in the “reaction notes” using the CellDesigner software. The network was based on 24 unique articles that are present in the PubMed database. In the PD network, genes and their corresponding protein (coloured-code boxes) were present that could be associated with ATP- and DNA-binding, transcription function and gene products that could be associated with membrane and mitochondrion (Figure 2A). Other DE genes and their proteins were included as well. The grey-coloured genes and proteins were retrieved from the literature and formed together with the DE genes and their products a biochemical reaction network that represented molecular mechanisms in the thyroid. To this network gene expression data was applied (colour coded genes and their products) which allowed better understanding of the effect of KI on gene expression in the network. Certain biological processes (ochre coloured hexagons) including the AKT pathway, Notch pathway and apoptosis are modulated by the network (Figure 2C)

### Network inference for the cortex upon repetitive KI treatment

For the cortex, we applied the same strategy. Transcriptomics showed 94 DE transcripts/genes and to those genes the following function can be associated: ATP-, DNA- and RNA-binding, transcription, neurogenesis and associated with the membrane (Figure 1B). The heat map showed a distinct difference in gene expression between the two groups.

The inferred network showed one highly connected network that is scale-free meaning there are few nodes that have many connections (Hubs) and there are many nodes with a few connections indicating this network is not a random network^22^. The hubs in this network were *hist3h2ba, ralgapa1, pik3ca and chmp4bl1*. Their molecular functions were DNA binding, GTPase activator, ATP binding, vacuolar transport, respectively (supplementary materials table 2). The genes *elavl2, gtpbp4, hmg1/1, hsp90aa1, katnal1, swi5*, and the predicted gene *cdkl5* were selected by sPLS-DA and were all under expressed in the ITB group. They were associated with nucleotide binding, GTPase activity, DNA-binding, neurogenesis, ATP-binding, DNA repair (supplementary materials table 2). The PD network was constructed with the aim of the inferred network by ShrinkNet and the selected genes by sPLS-DA. The data to support the biochemical reactions between the nodes within the PD network were obtained from published articles accessible by PubMed. The PD network from the cortex contained 130 chemical species including genes, RNA protein, anti-sense RNA molecules and 92 edges/biochemical reactions that were annotated within the “reactions note” (Figure 3C). The network was based on 44 unique articles that were present in the PubMed database. In the PD network, genes and their corresponding protein (coloured-code boxes) were present that could be associated with ATP- and DNA-binding, transcription function and gene products that were associated with membrane, mitochondrion and neurogenesis (Figure 3A). Other DE genes and their proteins were included as well. The grey-coloured genes and proteins were retrieved from the literature and together with the DE genes and their gene products they formed a biochemical reaction network that represented molecular mechanisms in the cortex. To this network gene expression data was applied (colour coded genes and their products) which allowed better understanding the effect of KI on gene expression in the network. Certain DE genes including katnal1, hmgb1, cacna1a and ralgapa1 modulate the biological processes (Ochre coloured hexagons) associated with memory, intelligence, learning, neurite outgrowth and branching, transmitter release and brain development (Figure 3C).

**Figure 3.**
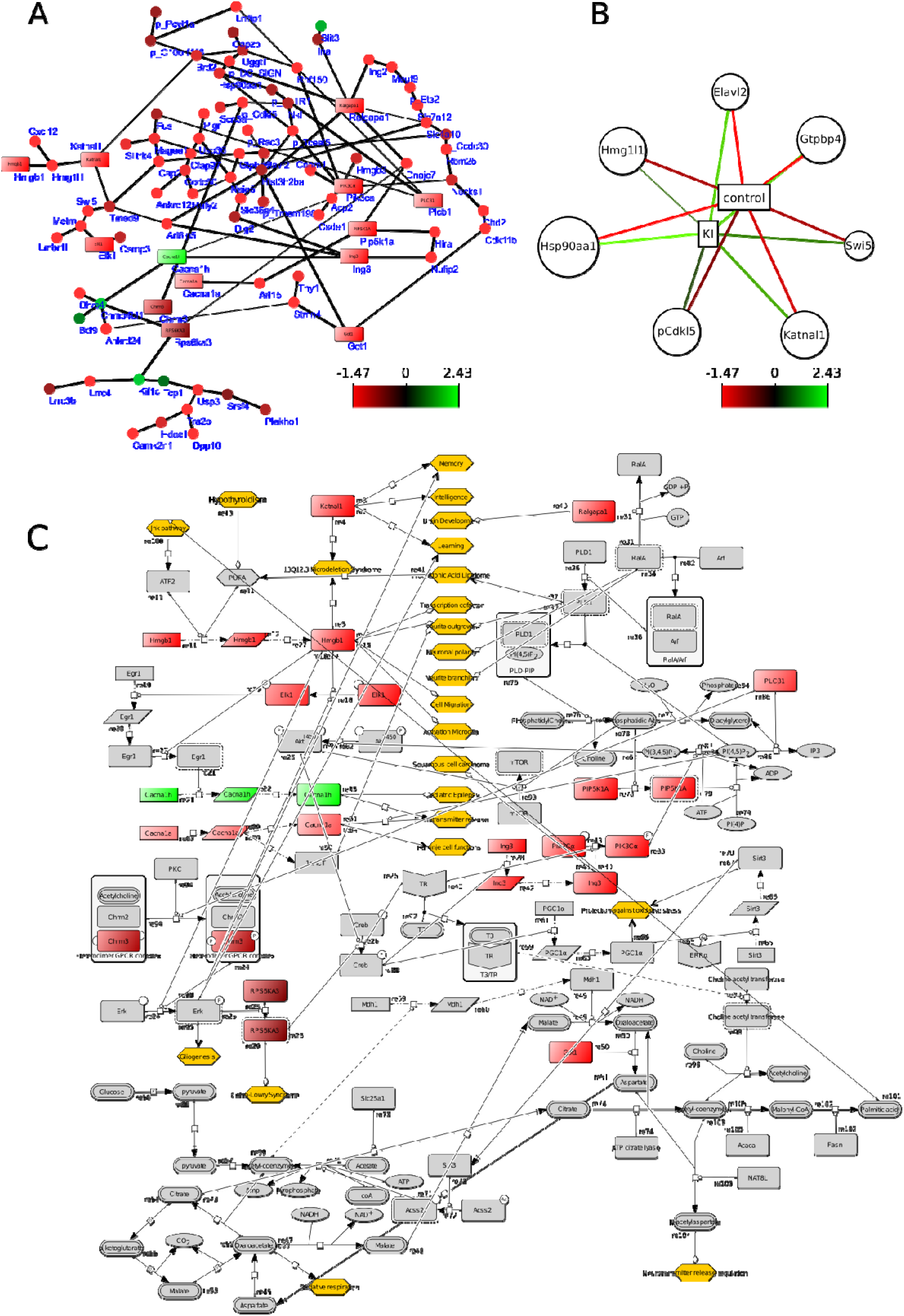
Selected diagrams showing correlations between genes from the cortex. The colour of the nodes indicates up or down regulation. Ochre coloured hexagons show phenotypes/biological processes modulated by the network. Figure 3A shows a undirected network of DE genes using Bayesian simultaneous-equation model with global-local shrinkage^12^. The nodes (closed circles and squares) represent genes and edges represent correlation between genes. The square nodes are genes that can be found in the process descriptive network as well (figure 2C). Figure 2B shows the results of sPLS-DA: genes that contribute most to either the control group or the IK exposed group, respectively. These genes will be used as a starting point for constructing the PD network (figure 2C). Figure 2C shows a process descriptive network that represents molecular mechanism^10^. Squares that are colour coded shows the fold changes of DE genes. The grey boxes are molecules obtained from published articles. All edges have been described before in the scientific article.

### Between-organ integrative network: thyroid and cortex upon repetitive KI administration

It was shown before that thyroid hormones are crucial for brain development^23^, we determined if gene expression of the two organs (thyroid and cortex) were correlated, Bayesian statistics using ShrinkNet and DIABLO framework using mixOmics were applied on genes from both organs. The ShrinkNet inferred network showed both “between” and “within” organs correlations (Figure 4A) and the inferred network obtained by DIABLO connected the selected genes in both organs which maximize the “between-organ” transcriptomic expression correlation (Figure 4B).

**Figure 4A.**
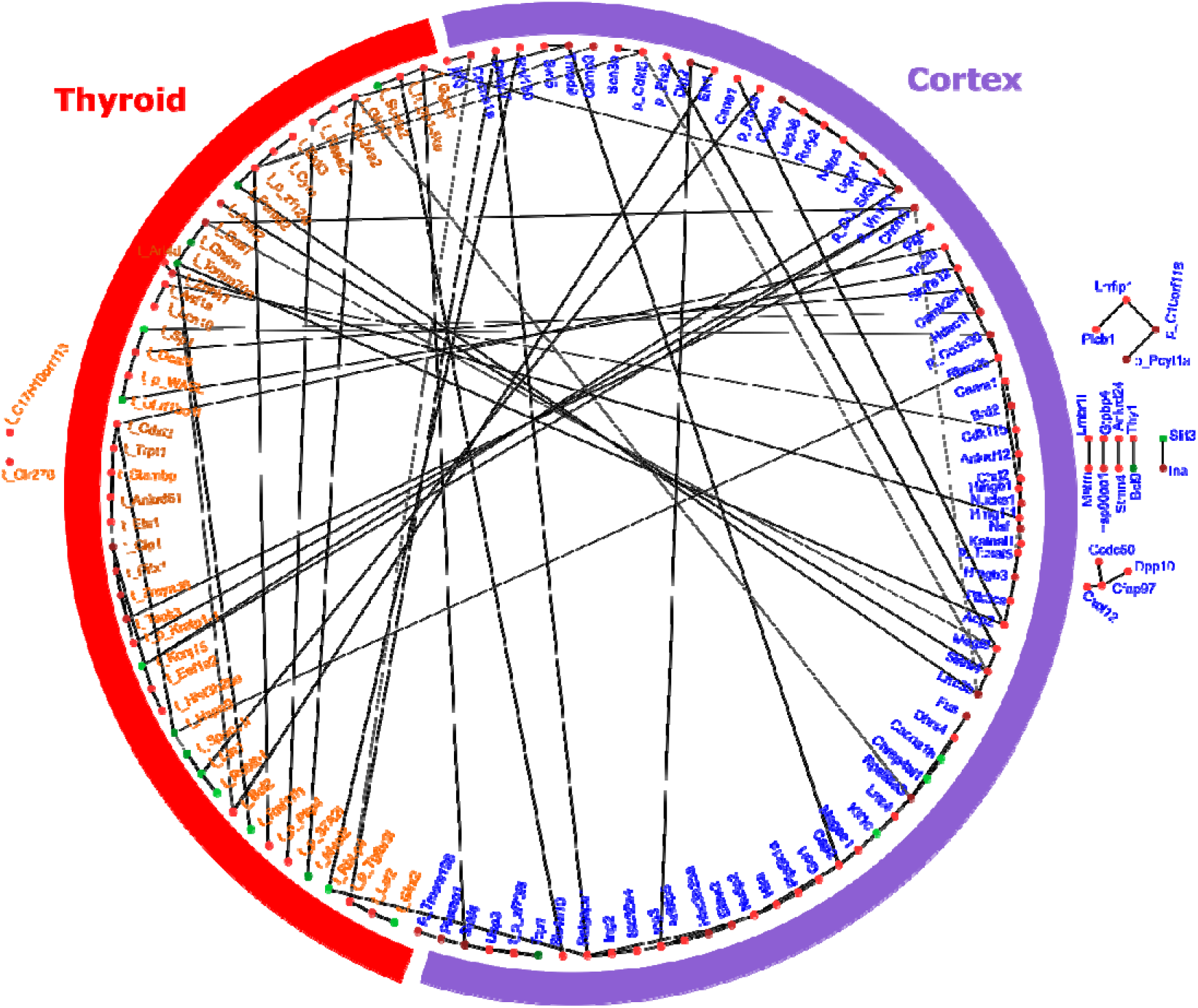
ShrinkNet network of DE genes obtained from thyroid and cortex respectively. There are edges present that show correlation between expression of genes within the same organ (thyroid or cortex) but there are also edges present showing correlation between genes from both organs. In addition, there are genes present from one organ that either do not have edges to genes of the other organ or do not have edges to genes that have edges to genes of the other organ (nodes that are outside the circle).

**Figure 4B.**
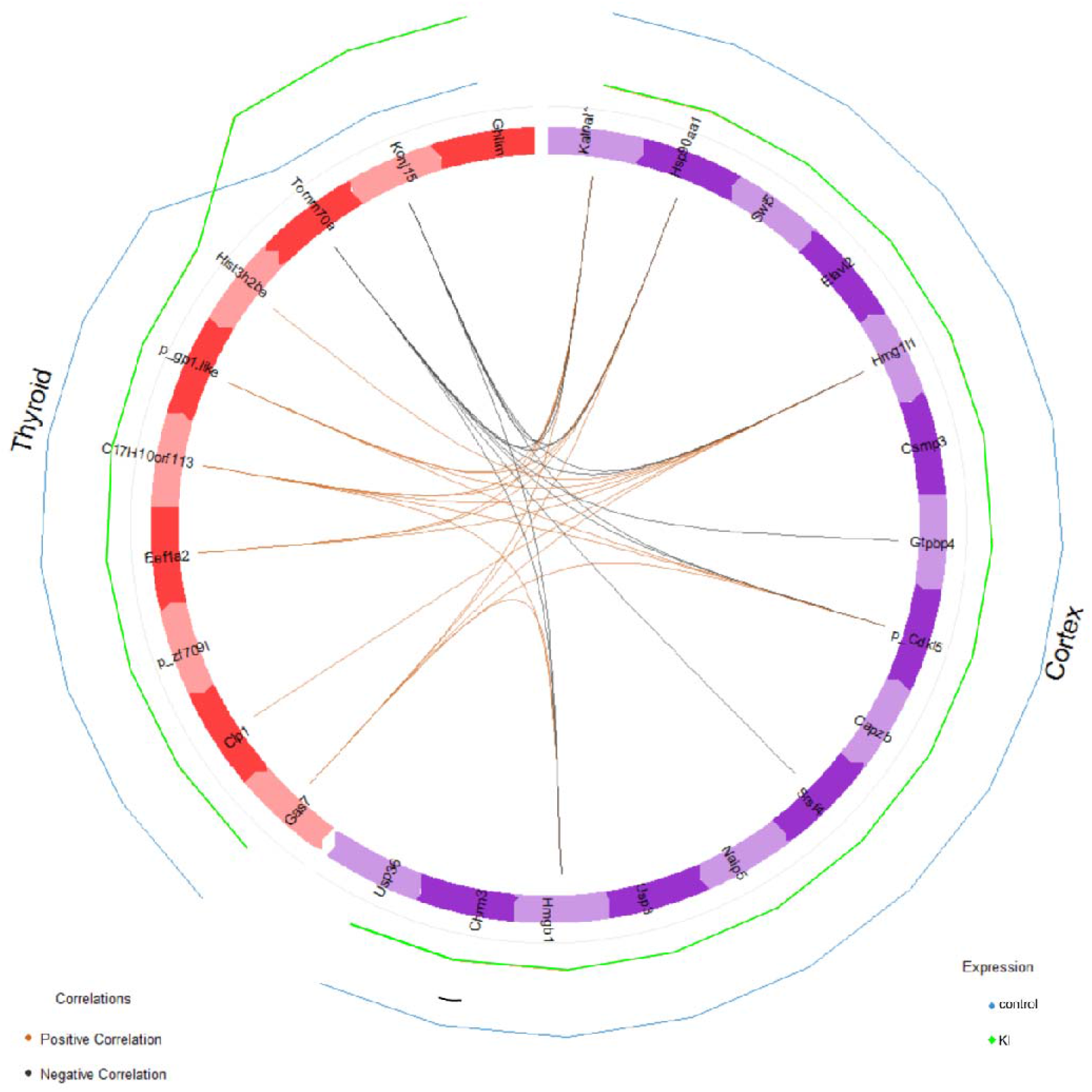
The DIABLO inferred gene network is represented as a circle where the red part of the circle represents the thyroid with genes that contribute strongly to the separation of the control group and the KI exposed group. The purple part of the circle represents the cortex with genes that contribute strongly to the separation of the KI group and the control group. The edges show correlation (with a cut-off of r=0.82) between genes that can be either negative (black colour) or positive (orange colour). These genes can be used as a starting point for creating the PD network (figure 4C).

To “validate” the results of the ShrinkNet and DIABLO frameworks, a PD network was constructed (Figure 4C). The networks showed a process descriptive diagram constituted of genes that were differential regulated in either the cortex or thyroid. One process that was of particular interest was the activation of Chrm2/3 complex by acetylcholine that lead to activation of the ERK transcription factor (TF) which in turns increased transcription of the Sox9 TF. Sox9 induced the transcription of *gas7* and the GAS7 protein with the WASL protein (complex) affected the neuronal outgrowth. The gene *chrm3* and the genes *gas7* and *wasl* were differential expressed in cortex and thyroid, respectively. In addition, the genes *tomm70a, bcl2*, and *hsp90aa1, hmgb1* from thyroid and cortex, respectively, were implicated in mitochondrial function and mitophagy.

**Figure 4C.**
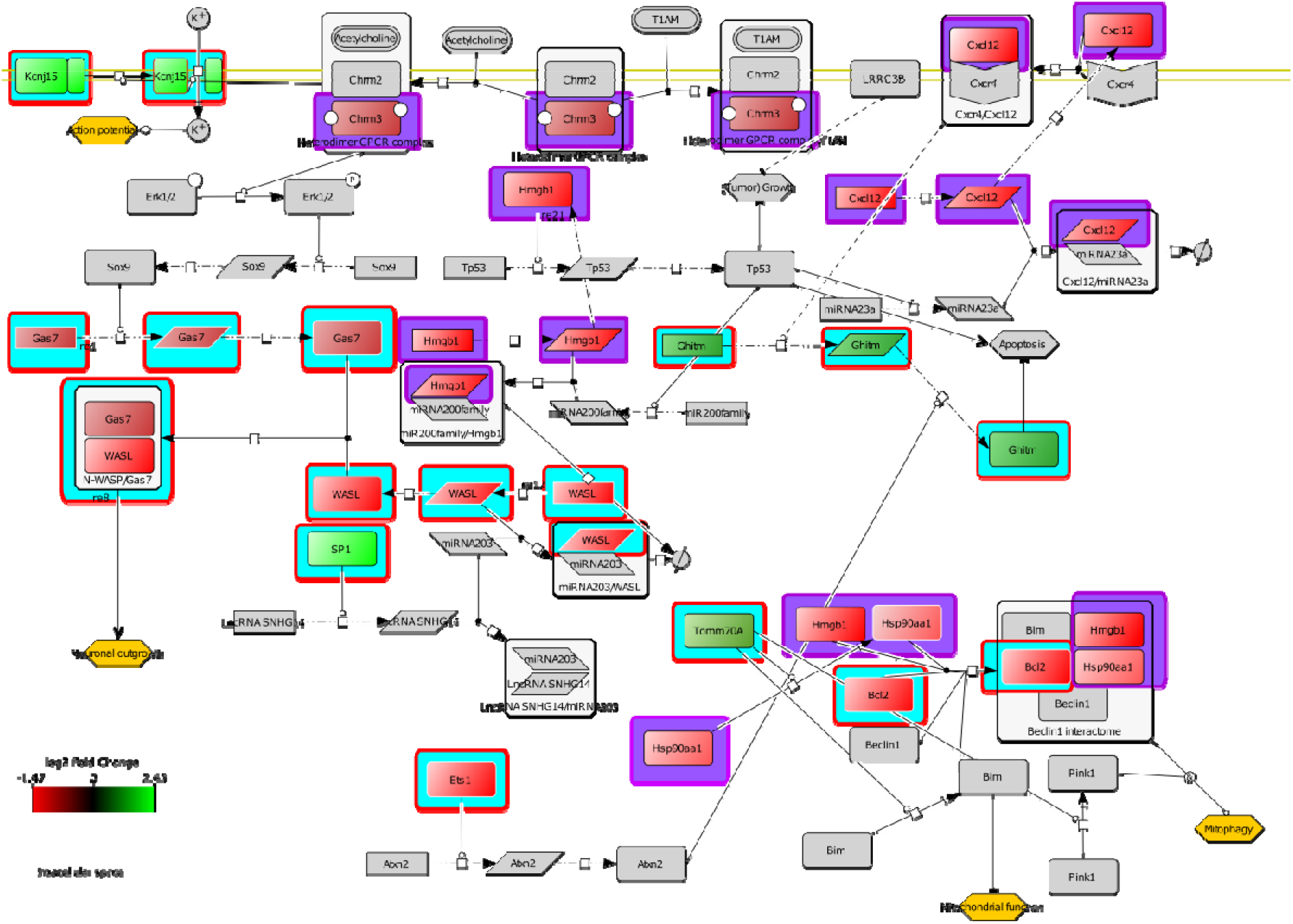
A process descriptive network shows DE genes that are obtained from the thyroid (light blue background) and cortex (purple background), respectively. The information about the interaction between these genes and their corresponding proteins and the interaction with other molecules is obtained from published articles. This figure demonstrates molecular mechanisms between genes from two different organs that impact neuronal outgrowth, and action potential in neurons and mitochondrial function and mitophagy (Ochre coloured hexagons) within the cell.

## 4. DISCUSSION

The Fukushima disaster with repetitive release of radionucleotides in the atmosphere has revealed the limitation of a strategy based on a single ITB may not be sufficient to protect the thyroid. The established doctrine of single ITB needs to be evaluated in case of repetitive releases of radionucleotides and indeed repetitive or multiple doses of ITB protects against prolonged exposure of ^131^I in adults^24^. However, for higher risk groups *i.e*. pregnant women and unborn children, a negative impact may be expected such as hypothyroidism and related CNS impairment, decrease in TSH and T4 and change in behaviour were noted^8,25–28^. The molecular mechanisms that drive these AO are currently unknown and need further investigation to propose adapted/optimised KI administration. The application of systems biology may provide additional insights on the effect of KI and may be able to propose a molecular mechanism^29–31^.

In this study we have focussed on the thyroid and cortex as these organs are the major target organs during embryonal development^32,33^. Indeed, for the cortex and thyroid, distinct gene expression profiles between the control and KI-treated groups have been observed even at 51 days after birth, while in adult rat no difference in gene expression between the control and KI-treated groups have been observed^6^. We have hypothesized that the observed late gene expression profile (Figure 1) is a result of KI-disturbed foetal development. Actually, it has been shown before that treatment of the mother can affect the progeny during pregnancy^34–37^.

The gene expression profile for the cortex has shown that DE genes are associated with DNA binding: *hmgb1, hmgb3, hmg1/1, ing2, csde1, lrrfip1, hist3h2ba*; transcription: *brd2, ing3, elk1, hdac1l, lrrfip1*; and neurogenesis: *ina, metrn, slit3*. In the thyroid, DE genes are associated with transcription: *sp1, ets1, pitx1*; DNA binding: *sp1, ets1, hist3h2ba, pitx1*; as membrane components: *stambp, olr278, kcnj15, extl3, lat2, ghitm, slc24a2, bcl2, glra2, gabbr1*. Certain of these genes, identified in the above functional groups, are important in the brain development^38–41^. However, to show how the above listed genes are interconnected, a network construction based on (DE) genes will help in understanding cellular processes that may be deregulated during ITB.

Indeed, network construction is a well-established method to asses patho-physiology^42^ and can provide new insights in the etiology of diseases^31^.

We have taken advantage of different methods for constructing a network i.e. ShrinkNet and sPLS-DA/Diablo. For the thyroid, the four sub-networks may represent four different phenotypes that may contribute to the same dysfunction. The network of the cortex shows one network with few hubs (nodes with many connections). Hubs are essential nodes and targeting these nodes impacts the network significantly while non-hub nodes are less important and mutations of those node have less impact on the network^22^. Due to its complexity (many more connections compared to the thyroid network) it is more difficult to find genes that may lead to synthetic lethality^43^ and therefore this network is more robust against deregulation of homeostasis than the thyroid one. Because the thyroid network is less robust, it is easier to modulate this network e.g. a low dosage KI may have a major impact on thyroid hormone synthesis and this has in turn a major effect on the development of the cortex^44^.

The obtained networks show connections between genes in terms of statistical correlations. The CellDesigner software permits to add a plausible and biochemical interpretation to the inferred network. The PD network of the thyroid has shown that the observed DE genes may affect apoptosis although it is not clear if this is either positive or negative as Tp53 (apoptosis activator) is inhibited by the genes *pitx1, rbpp4* and *sp1* while taking their expression profile into account. On the other hand, the PI3K/AKT pathway and *bcl2* are inhibited which are known as apoptosis inhibitors^45,46^. The PD network suggests that apoptosis is a target of repetitive ITB; KI-induced hypothyroidism can activate apoptosis through *bcl2*. This result was unexpected after the initial DE genes approach since no apoptotic function arose among the listed cellular processes. It has been demonstrated before that the excess of KI provoked the activation of the molecular mechanisms associated with apoptosis^47,48^. Furthermore, AKT has been found to be important in the development of the brain upon iodine deficiency-induced hypothyroidism^49^. In the cortex, the genes *hmgb1* and *katnal1* have many direct arches with phenotypes that are associated with memory, learning, brain development, etc. These direct connections have been described before but the exact mechanisms behind these observations are unknown. In addition, the up-regulated gene *cacna1h* is involved in childhood absence epilepsy^50^ and the down-regulated gene *cacna1a* is involved in the activation of muscarinic M1-class receptors and neurite outgrowth^51,52^, and the down-regulated *got1* gene may be negatively impact the synthesis of acetylcholine. The PD diagram shows putative mechanisms how these DE genes may impact the brain development and function. Indeed, Lebsir and colleagues have shown that multiple administration of KI during pregnancy lead to alterations in behaviour in the progeny of rats^8,9^.

Thyroid and cortex developments are connected during fetal development. Repetitive ITB occurring this period leads to joined impact on these two organs. This motivated to connect the DE genes from both the cortex and the thyroid into one network which suggests: (i) cortex to thyroid connection through chrm2 to *gas7*; (ii) thyroid to cortex through tomm70a which orchestrate complex formation that drives apoptosis/mitophagy (Figure 5). Mitophagy is implicated in neurodegenerative diseases including Alzheimer and Parkinson^53^. Connection (i) is associated with acetylcholine and neurite outgrowth. This is in accordance with our hypothesis and other studies^9,54–56^ that transient hypothyroidism impact acetylcholine pathway and the apoptotic pathway. These directed biochemical reactions contrast with the non-directed statistical inferred networks by improving the mechanistic understanding of the integrated response to repetitive ITB.

**Figure 5.**
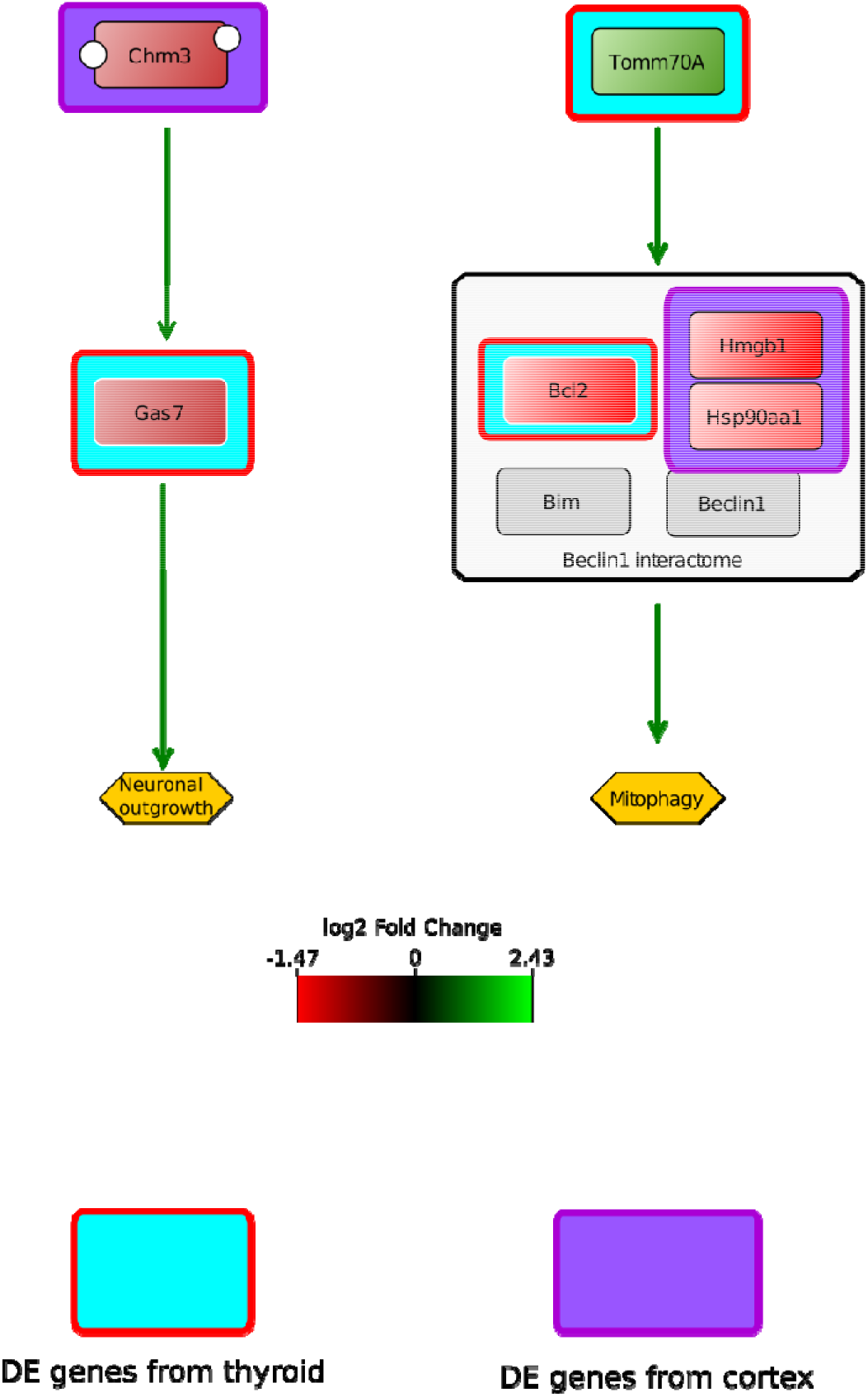
A simplified diagram derived from figure 4 showing the positive effect of DE genes that have been identified using the sPLS-DA/DIABLO platform on neuronal outgrowth or mitophagy. Their effect is based on based on their involvement in the biochemical reaction network (PD diagram, Figure 4). The gene expression is colour coded and added to the genes. In case of the genes *chrm3* and *gas7*, both are under-expressed and this may result in diminished neuronal outgrowth as during homeostatic conditions both genes have an activating effect on neuronal outgrowth. The gene *tomm70a* is over-expressed and this gene has a positive effect on the formations of the Beclin1 interactome which may have an activing effect on mitophagy during homeostatic conditions. However, three DE genes are under-expressed and this may result in no interactome formation. Or *tomm70a* is over-expressed to compensate for the loss of the Beclin1 interactome.

### Final conclusion

Our vigorous work flow is to our knowledge the first that combines advanced statistical network inference approaches with biochemical network validation which establish a thyroid/cortex integrated molecular mechanism between acetylcholine, neurogenesis, mitophagy in a repetitive ITB *in utero* framework. This study may give new insights for improvement of knowledge of adverse effect of ITB for the protection of pregnant women.

## ACKNOWLEDGEMENTS

This study was part of the PRIODAC research program funded by ANR-RSNR: ANR-11-RSNR-0019-01.

## 6. SUPPLEMENTAIRY INFORMATION

Suppl Table1. Differentially expressed genes in the cortex

Suppl table2. Differentially expressed thyroid

## REFERENCES

1. Ron, E. Thyroid cancer incidence among people living in areas contaminated by radiation from the Chernobyl accident. Health Phys. 93, 502–11 (2007).

2. Furukawa, K. et al. Long-term trend of thyroid cancer risk among Japanese atomic-bomb survivors: 60 years after exposure. Int. J. cancer 132, 1222–6 (2013).

3. WHO. Iodine thyroid blocking: Guidelines for use in planning for and responding to radiological and nuclear emergencies. (2017).

4. Dreger, S., Pfinder, M., Christianson, L., Lhachimi, S. K. & Zeeb, H. The effects of iodine blocking following nuclear accidents on thyroid cancer, hypothyroidism, and benign thyroid nodules: Design of a systematic review. Syst. Rev. 4, (2015).

5. Benderitter, M. et al. Do Multiple Administrations Of Stable Iodine Protect Population Chronically Exposed To Radioactive Iodine: What Is Priodac Research Program (2014-22) Teaching Us? Radiat. Prot. Dosimetry 182, 67–79 (2018).

6. Lebsir, D. et al. Effects of repeated potassium iodide administration on genes involved in synthesis and secretion of thyroid hormone in adult male rat. Mol. Cell. Endocrinol. 474, 119–126 (2018).

7. Lebsir, D. et al. Toxicology of Repeated Iodine Thyroid Blocking in Adult Rat. J. Pharm. Res. 3, (2018).

8. Lebsir, D. et al. Repeated potassium iodide exposure during pregnancy impairs progeny’s brain development. Neuroscience 406, 606–616 (2019).

9. Rosique, C. et al. Assessment of the effects of repeated doses of potassium iodide intake during pregnancy on male and female rat offspring using metabolomics and lipidomics. J. Toxicol. Environ. Health. A 1–13 (2019). doi: 10.1080/15287394.2019.1625474

10. Le Novère, N. Quantitative and logic modelling of molecular and gene networks. Nat. Rev. Genet. 16,146–58 (2015).

11. Ritchie, M. E. et al. limma powers differential expression analyses for RNA-sequencing and microarray studies. Nucleic Acids Res. 43, e47(2015).

12. Leday, G. G. R. et al. Gene Network Reconstruction using Global-Local Shrinkage Priors. Ann. Appl. Stat. 11, 41–68 (2017).

13. Rohart, F., Gautier, B., Singh, A. & Lê Cao, K.-A. mixOmics: An R package for ’omics feature selection and multiple data integration. PLoS Comput. Biol. 13, e1005752 (2017).

14. Lê Cao, K.-A., Boitard, S. & Besse, P. Sparse PLS discriminant analysis: biologically relevant feature selection and graphical displays for multiclass problems. BMC Bioinformatics 12, 253 (2011).

15. Singh, A. et al. DIABLO: an integrative approach for identifying key molecular drivers from multi-omic assays. Bioinformatics (2019). doi:10.1093/bioinformatics/bty1054

16. Shannon, P. et al. Cytoscape: A software Environment for integrated models of biomolecular interaction networks. Genome Res. 13, 2498–2504 (2003).

17. Funahashi, A., Morohashi, M., Kitano, H. & Tanimura, N. CellDesigner: a process diagram editor for gene-regulatory and biochemical networks. Biosilico 1, 159–162 (2003).

18. Hucka, M. et al. The systems biology markup language (SBML): A medium for representation and exchange of biochemical network models. Bioinformatics 19, 524–531 (2003).

19. Kitano, H., Funahashi, A., Matsuoka, Y. & Oda, K. Using process diagrams for the graphical representation of biological networks. Nat. Biotechnol. 23, 961–6 (2005).

20. Le Novère, N. et al. The Systems Biology Graphical Notation. Nat. Biotechnol. 27, 735–41 (2009).

21. Bonnet, E. et al. BiNoM 2.0, a Cytoscape plugin for accessing and analyzing pathways using standard systems biology formats. BMC Syst. Biol. 7, 18 (2013).

22. Barabási, A.-L. & Bonabeau, E. Scale-free networks. Sci. Am. 288, 60–9 (2003).

23. López-Espíndola, D.et al. Thyroid hormone availability in the human fetal brain: novel entry pathways and role of radial glia. Brain Struct. Funct. (2019). doi:10.1007/s00429-019-01896-8

24. Sternthal, E. et al. Suppression of thyroid radioiodine uptake by various doses of stable iodide. N. Engl. J. Med. 303, 1083–8 (1980).

25. Noteboom, J. L. et al. Protection of the maternal and fetal thyroid from radioactive contamination by the administration of stable iodide during pregnancy. An experimental evaluation in chimpanzees. Radiat. Res. 147, 691–7 (1997).

26. Shi, X. et al. Optimal and safe upper limits of iodine intake for early pregnancy in iodine-sufficient regions: a cross-sectional study of 7190 pregnant women in China. J. Clin. Endocrinol. Metab. 100, 1630–8 (2015).

27. Pearce, E. N., Lazarus, J. H., Moreno-Reyes, R. & Zimmermann, M. B. Consequences of iodine deficiency and excess in pregnant women: an overview of current knowns and unknowns. Am. J. Clin. Nutr. 104 Suppl, 918S–23S (2016).

28. Min, H. et al. Effects of Maternal Marginal Iodine Deficiency on Dendritic Morphology in the Hippocampal CA1 Pyramidal Neurons in Rat Offspring. Neuromolecular Med. 18, 203–15 (2016).

29. Lin, M., Ye, M., Zhou, J., Wang, Z. P. & Zhu, X. Recent Advances on the Molecular Mechanism of Cervical Carcinogenesis Based on Systems Biology Technologies. Comput. Struct. Biotechnol. J. 17, 241–250 (2019).

30. Capriotti, E., Ozturk, K. & Carter, H. Integrating molecular networks with genetic variant interpretation for precision medicine. Wiley interdiscip. Rev. Syst. Biol. Med. 11, e1443 (2019).

31. Wang, Z.-T., Tan, C.-C., Tan, L. & Yu, J.-T. Systems biology and gene networks in Alzheimer’s disease. Neurosci. Biobehav. Rev. 96, 31–44 (2019).

32. Agirman, G., Broix, L. & Nguyen, L. Cerebral cortex development: an outside-in perspective. FEBS Lett. 591, 3978–3992 (2017).

33. Fagman, H. & Nilsson, M. Morphogenesis of the thyroid gland. Mol. Cell. Endocrinol. 323, 35–54 (2010).

34. Dowling, A. L. & Zoeller, R. T. Thyroid hormone of maternal origin regulates the expression of RC3/neurogranin mRNA in the fetal rat brain. Brain Res. Mol. Brain Res. 82, 126–32 (2000).

35. Dong, J., Liu, W., Wang, Y., Xi, Q. & Chen, J. Hypothyroidism following developmental iodine deficiency reduces hippocampal neurogranin, CaMK II and calmodulin and elevates calcineurin in lactational rats. Int. J. Dev. Neurosci. 28, 589–96 (2010).

36. Sawano, E., Takahashi, M., Negishi, T. & Tashiro, T. Thyroid hormone-dependent development of the GABAergic pre- and post-synaptic components in the rat hippocampus. Int. J. Dev. Neurosci. 31, 751–61 (2013).

37. Kobayashi, K. et al. Perinatal exposure to PTU decreases expression of Arc, Homer 1, Egr 1 and Kcna 1 in the rat cerebral cortex and hippocampus. Brain Res. 1264, 24–32 (2009).

38. Yuan, A. & Nixon, R. A. Specialized roles of neurofilament proteins in synapses: Relevance to neuropsychiatric disorders. Brain Res. Bull. 126, 334–346 (2016).

39. Zhang, B. et al. Repulsive axon guidance molecule Slit3 is a novel angiogenic factor. Blood 114, 4300–9 (2009).

40. Wang, Z. et al. Meteorin is a chemokinetic factor in neuroblast migration and promotes stroke-induced striatal neurogenesis. J. Cereb. Blood Flow Metab. 32, 387–98(2012).

41. Angelopoulou, E., Piperi, C., Adamopoulos, C. & Papavassiliou, A. G. Pivotal role of high-mobility group box 1 (HMGB1) signaling pathways in glioma development and progression. J.Mol. Med. (Berl). 94, 867–74 (2016).

42. Hanash, S., Schliekelman, M., Zhang, Q. & Taguchi, A. Integration of proteomics into systems biology of cancer. Wiley Interdiscip. Rev. Syst. Biol. Med. 4, 327–37

43. Zhu, X., Gerstein, M. & Snyder, M. Getting connected: analysis and principles of biological networks. Genes Dev. 21,1010–24 (2007).

44. Bernal, J. Thyroid hormone regulated genes in cerebral cortex development. J. Endocrinol. 232, R83–R97 (2017).

45. Zhao, S. et al. Inhibitor of growth 3 induces cell death by regulating cell proliferation, apoptosis and cell cycle arrest by blocking the PI3K/AKT pathway. Cancer Gene Ther. 25, 2440–247 (2018).

46. Pentimalli, F., Grelli, S., Di Daniele, N., Melino, G. & Amelio, I. Cell death pathologies: targeting death pathways and the immune system for cancer therapy. Genes Immun. (2018). doi:10.1038/s41435-018-0052-x

47. Basalaeva, N. L., Sychugov, G. V, Strizhikov, V. K. & Mikhailova, E. N. Iodine concentration and signs of apoptosis in the thyroid and pituitary of female rats after different single doses of potassium iodide. Endocr. Regul. 45,183–90 (2011).

48. Zhang, M. et al. Effect of Excessive Potassium Iodide on Rat Aorta Endothelial Cells. Biol. Trace Elem. Res. 166, 201–9 (2015).

49. Wang, Y. et al. Neurotoxicity of developmental hypothyroxinemia and hypothyroidism in rats: Impairments of long-term potentiation are mediated by phosphatidylinositol 3-kinase signaling pathway. Toxicol. Appl. Pharmacol. 271, 257–65 (2013).

50. Wang, G. et al. Ca V 3.2 calcium channels control NMDA receptor-mediated transmission: a new mechanism for absence epilepsy. Genes Dev. 29, 1535–1551 (2015).

51. Du, X. et al. Second cistron in CACNA1A gene encodes a transcription factor mediating cerebellar development and SCA6. Cell 154, 118–33 (2013).

52. Perez-Burgos, A., Prieto, G. A., Galarraga, E. & Bargas, J. CaV2.1 channels are modulated by muscarinic M1receptors through phosphoinositide hydrolysis in neostriatal neurons. Neuroscience 165, 293–299 (2010).

53. Ding, W.-X. & Yin, X.-M. Mitophagy: mechanisms, pathophysiological roles, and analysis. Biol. Chem. 393, 547–64 (2012).

54. Zhang, L. et al. Effect of maternal excessive iodine intake on neurodevelopment and cognitive function in rat offspring. BMC Neurosci. 13, 121 (2012).

55. Serrano-Nascimento, C., Salgueiro, R. B., Pantaleão, T., Corrêa da Costa, V. M. & Nunes, M. T. Maternal Exposure to Iodine Excess Throughout Pregnancy and Lactation Induces Hypothyroidism in Adult Male Rat Offspring. Sci. Rep. 7, 15591 (2017).

56. Serrano-Nascimento, C. et al. Iodine excess exposure during pregnancy and lactation impairs maternal thyroid function in rats. Endocr. Connect. 6, 510–521 (2017).

